# A conserved role of α2δ subunit of calcium channel in nicotine motivated behavior

**DOI:** 10.1101/2022.06.21.497113

**Authors:** Chinnu Salim, Enkhzul Batsaikhan, Ann Ke Kan, Hao Chen, Changhoon Jee

**Affiliations:** Dept. of Pharmacology, Addiction Science and Toxicology, College of medicine, University of Tennessee Health Science Center, 71 S Manassas St., Memphis, TN

**Keywords:** Nicotine, motivated behavior, CCP, *C. elegans*, α2δ, nAChRs

## Abstract

Identifying genetic variants associated with nicotine-motivated behavioral traits is an important strategy to understand the fundamental mechanisms underpinning smoking and tobacco abuse. For suitable emulation of behavioral phenotype with the full advantage of this invertebrate model, we newly established a worm model of nicotine seeking by Conditioned Cue Preference (CCP). We demonstrated that *C. elegans* also exhibited pivotal features of nicotine-motivated behaviors as in mammals. First, we identified the nicotine-elicited cue preference is mediated by nicotinic acetylcholine receptors in worms. Additionally, we exhibited dopamine is also required for the development of CCP. Subsequently, we identified the nAChRs subunits associated with the facilitation of nicotine preference. Accordingly, we validated human GWAS candidates associated with nicotine dependence involved in the role of those nAChR subunits. we addressed the cross-species functional validation to determine the GWAS candidate genes have authentic roles in nicotine seeking associated with tobacco abuse. The loss of function strain of *CACNA2D3* orthologue, calcium voltage-gated channel auxiliary subunit alpha2delta 3, was tested for CCP. We also tested the knock-out (KO) strain of the *CACNA2D2* orthologue, calcium voltage-gated channel auxiliary subunit alpha2delta 2, which is closely related to *CACNA2D3* in the same family and shared the human smoking phenotypes. Our orthogonal test suggests the functional conservation of the α2δ subunit of calcium channel in nicotine motivated behavior.

## Introduction

Tobacco abuse has been a major public health concern and smoking is still a leading cause of preventable death ^1^. Tobacco abuse is considerably heritable. ^2-6^. Nicotine dependence has been considered a hallmark in the progress and maintenance of tobacco abuse and human population genetics has identified statistically significant gene variants relevant to nicotine dependence. ^7-9^. Thus, the identification of genetic mechanisms underlying behavioral traits is an important strategy for understanding the underpinning mechanism of nicotine dependence. Although genome-wide association studies (GWAS) have been successfully identified numerous Single Nucleotide Polymorphisms (SNPs) associated with substance use disorder (SUD) over the past decade ^9-14^, most of the candidate genetic variants have not been independently validated or improved our understanding of nicotine dependence.

We, therefore, exploit the rapid genetic workflow of *C. elegans*, which has a simple nervous system but completely defined connectome^15-17^, as a tool for accelerating functional validation of GWAS candidates associated with smoking/nicotine self-administration behavior. *C. elegans* responds to abused substances in a way that mimics substance-dependent behaviors observed in mammals ^18-27^. Hence, worms have been a powerful model for SUD and nicotine dependence. *C. elegans* exhibit nicotine withdrawal-dependent behavior and state-dependent development of chemical preference, analogous to mammalian studies ^22,26,27^. Accordingly, we established nicotine Conditioned Cue Preference (CCP) to measure the nicotine preference and seeking in *C. elegans*. Here, we are demonstrating nicotine and withdrawal elicits CCP in *C. elegans*. The CCP assay stably elicits acquisition, progress, and extinction of nicotine-paired cue preference. The CCP assay also reveals CPP properties in mammals that are mediated by nicotinic acetylcholine receptors (nAChRs) and dopamine.

Subsequently, we tested nAChRs mutant animals in CCP to define nAChR subunits that specifically act on nicotine seeking. We then tested GWAS candidates of human smoking and nicotine dependence associated with the role of those nAChR subunits. Here, we suggest α2δ subunit of Voltage-Gated Calcium Channel (VGCC) is required for the nicotine preference affecting the progression of nicotine dependence.

## Results

### Establishment of CCP (Conditioned-Cue Preference)

Psychostimulants including nicotine elicit a CPP in rats and mice ^28-30^. We adapted the mammalian Conditioned Place Preference (CPP), a form of associative learning used to study the rewarding and aversive effects of drugs, to develop nicotine CCP and determine the nicotine preference and seeking in *C. elegans*. We have found that hexane, an alkane volatile odorant, is a neutral stimulus to *C. elegans* and is suitable as a conditioned stimulus (CS). Hexane was tested in numerous ranges of concentrations (Fig. 1a). *C. elegans* showed no preference at any tested concentrations (Hexane; 98.5% as non-diluted, 10^−1^, and 10^−2^ diluted). Subsequently, classical conditioning was performed using nicotine as an Unconditioned Stimulus (US) (Fig. 1b). The nicotine concentration and withdrawal time from nicotine were determined based on the behavioral and physiological response to various concentrations shown in previous studies ^23^.

**Fig.1.**
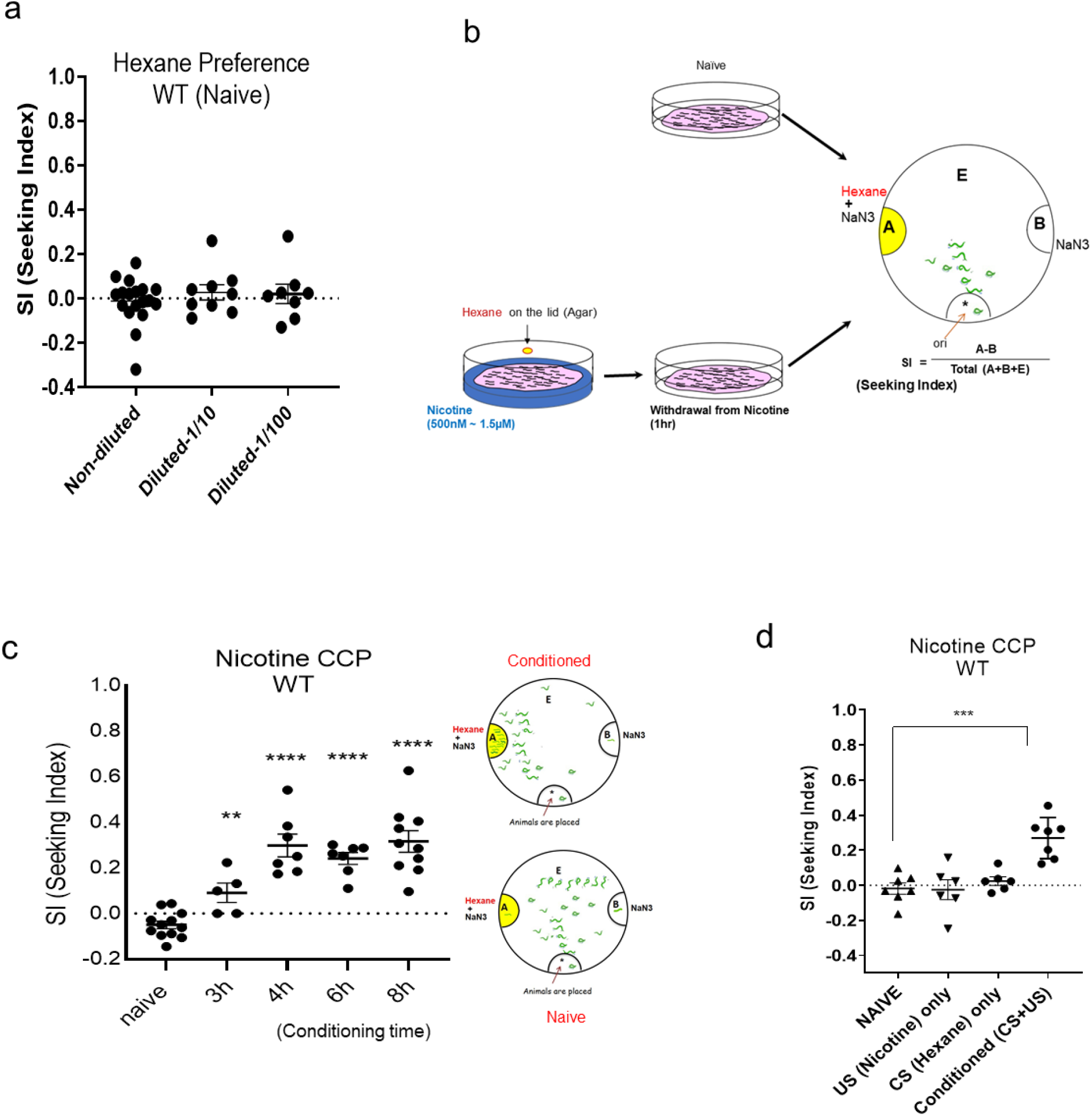
Nicotine Conditioned Cue Preference (CCP) using hexane as conditioned stimulus (CS). (**a**) Identification of hexane, as a neutral odor substance to naïve animals. One-way ANOVA of chemotaxis in wild-type animals to various concentrations of hexane did not show significant differences (p=0.6136, F(2, 32)=0.02649). (b) The diagram of nicotine Conditioned Cue Preference (CCP) using hexane as Conditioned Stimulus. 1-day adult Wild-type worms were pre-incubated with 1.5 µM nicotine and 2µl of non-diluted hexane for conditioning. The conditioned worms were transferred to OP50-Bacteria seeded plate then 1 hour later withdrawn worms from nicotine were moved to the chemotaxis assay plate. (c) Wild-type *C. elegans* develops CCP after chronic conditioning and following withdrawal from nicotine. (d) The CCP development by nicotine conditioning was validated by pretreatment of US only or CS only. US only; 4hr treatment of nicotine alone (1.5 µM), CS only; 4hr treatment of hexane alone, Conditioned (US + CS); 4hr conditioning of nicotine (1.5 µM) and hexane, all of those were withdrawn for 1 hour before chemotaxis to hexane. Each dot represents trial of population assay. (****, One-way ANOVA, F(4, 36)=1.683).

Worms first respond to the psychostimulant, nicotine by increasing motility at about 1.5 μM concentration (^26^), although, high concentrations (100 µM) of nicotine will induce locomotor paralysis in *C. elegans*; presumably due to acutely activating acetylcholine-sensitive ion channels on the worm’s motor neurons and muscles (^23^). Furthermore, worms show nicotine-induced motivated behavior over a range of concentrations (^31^). In addition, nicotine withdrawal causes locomotion stimulation in worms as a withdrawal symptom. The 1.5 μM concentration was sufficient to induce this nicotine-dependent stimulation of locomotion, thus we mainly used it for our CCP assays. Wild-type animals successfully develop acquisition of nicotine Conditioned Cue Preference (CCP) after a prolonged time of association in time-dependent manner (Fig.1c). In a similar manner, other concentrations of nicotine (which were higher but not paralyzing the worms) also successfully elicited the preference. Seeking Index (SI) is obtained as represented in Fig. 1b. A high SI indicates that the nicotine-paired cue acts as a strong attractant, which corresponds to the development of preference by the conditioning with reinforcing drug. Prolonged conditioning of CS (hexane) and US (nicotine) leads to the development of nicotine-paired cue preference, although hexane is a neutral olfactory stimulus to naive animals. The CCP was validated by pretreatment of US only, CS only, or Conditioned (CS+US), respectively. CCP was not developed by US only or CS only, whereas conditioning occurred and facilitated CCP when CS was paired with the US (Fig.1d).

We also demonstrate that CCP induced by nicotine is mediated by dopamine signaling. The development of CCP was impaired in the KO mutant animals of *cat-2*, tyrosine hydroxylase in *C. elegans*, which was dopamine deficient (Fig.2). It suggests that the CCP of worms has an evident face value, the recapitulation of the pivotal features of nicotine-dependent behaviors in mammals, in which nicotine-caused increased dopamine mediates nicotine-induced motivated behavior ^32, 33^. Additionally, wild-type animals can also represent the extinction of CCP, greatly reduced paired rewarding. Expression of nicotine-induced CCP was abolished in subsequent chemotaxis assays after the presentation of CS (hexane) alone in the absence of US (nicotine) during the withdrawal period (Fig.3a). Therefore, it feasibly suggests that CCP can be used to investigate the genes and pathways associated with reinstatement. Accordingly, to further investigate the underlying mechanism involved in the regulation of CCP in the neural circuits, we questioned neural circuits that mediate positive chemotaxis to CS that were previously neutral but acted as attractants after conditioning. Chemotaxis behaviors are regulated primarily by the chemosensory neurons and modulated by integration of signaling with interneurons ^34^,^35^. *C. elegans* have 32 presumed chemosensory neurons that detect a variety of olfactory and gustatory cues ^36-39^. In worms, AWC and AWA, ciliated chemosensory neurons, mediate attraction to the volatile odorants ^40^. We exploited AWC-ablated animals to test in CCP. Killing a pair of AWC neurons via expression of reconstituted Caspase^41,42^ resulted in impaired CS (hexane) preference after conditioning with US (nicotine) (Fig. 3b), indicating that the primary sensory neurons are AWC head neurons for attraction to hexane after conditioning.

**Fig.2.**
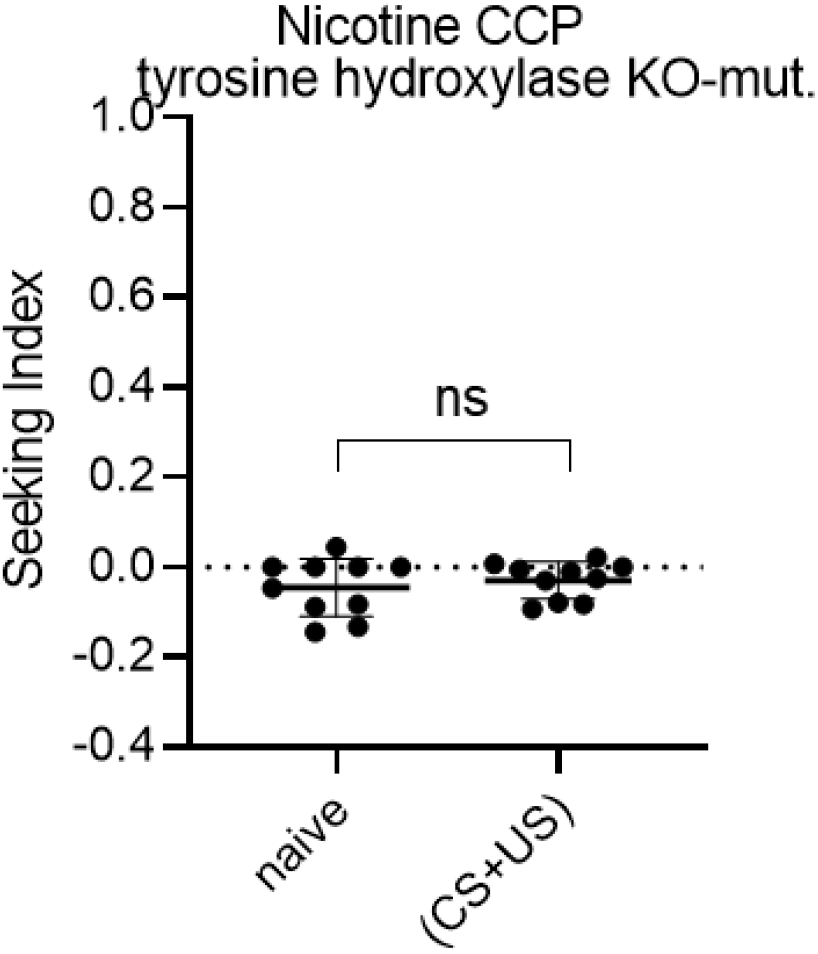
Dopamine is required to develop CCP. A *cat-2* encodes a tyrosine hydroxylase, which catalyzes the conversion of tyrosine to L-DOPA, the biosynthetic precursor of dopamine. Conditioned (US + CS); 4hr conditioning of nicotine (1.5 µM) and hexane, animals were withdrawn for 1 hour before chemotaxis to hexane. Each dot represents trial of population assay. P=0.7527(Mann-Whitney test).

**Fig. 3.**
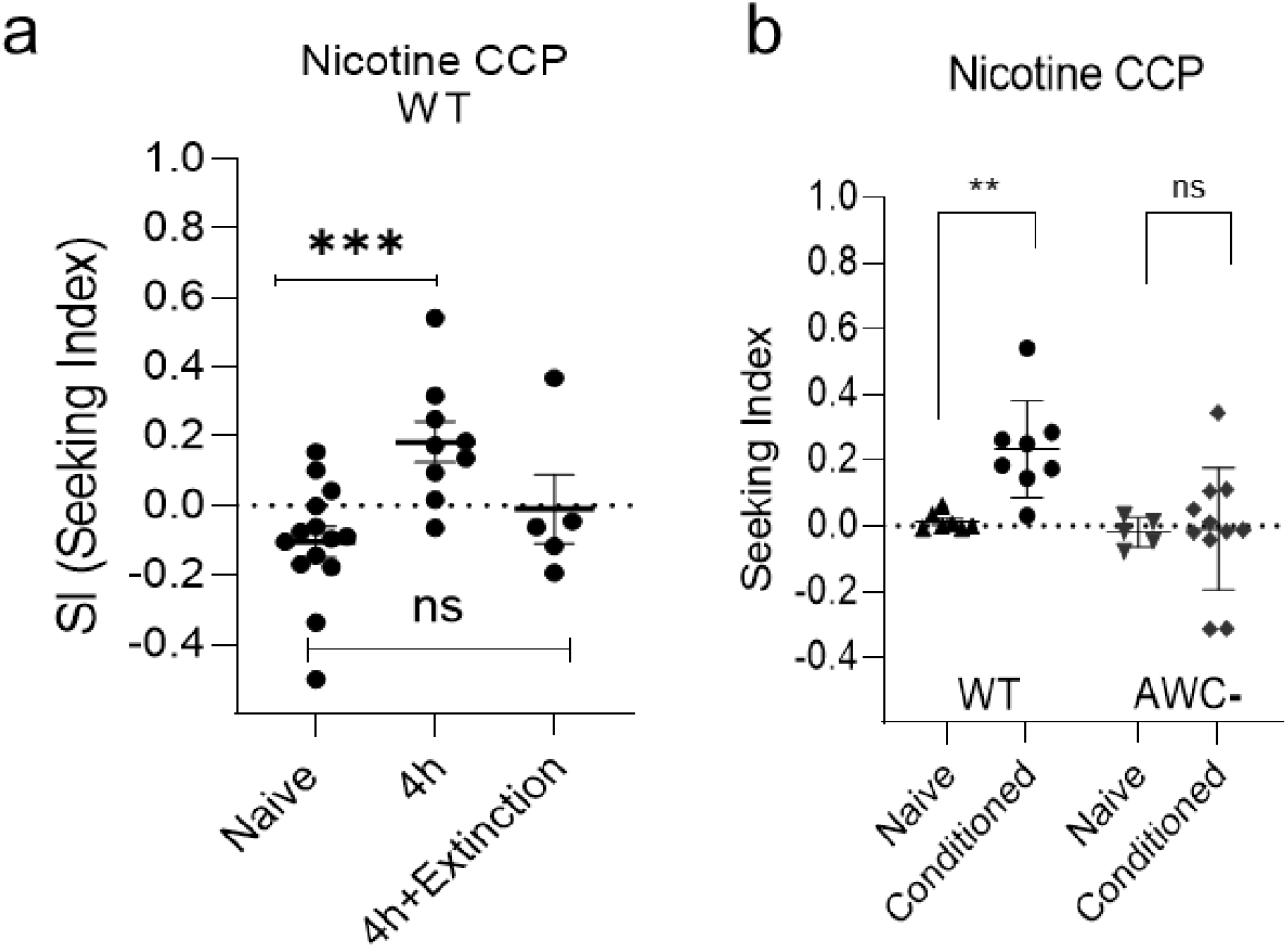
Characteristics of CCP. (a) Wild-Type *C. elegans* learns extinction of CCP. Each dot represents trial of population assay. **, P<0.01; ***, P<0.001 (Mann-Whitney test). (b) Nicotine Conditioned Cue Preference (CCP) of AWC neuron ablated animals. Single session of 4 hours CCP on 1.5 µM nicotine plates. Single session of 4 hours CCP on 1.5 µM nicotine plates

In the laboratory, *C. elegans* is reared in agar plates seeded with OP50 bacteria as a food source. Since cultivation without food has been used for odor/starvation conditioning paradigm for Conditioned Place Aversion (CPA), in order to dispel any controversy about the cultivation environment in the CCP assay, the nicotine conditioning and withdrawal process in CCP assay was conducted in the presence of OP50 bacteria on nicotine plates (Fig. 1, 2 and 3). However, we expanded the usage of the CCP to confirm the reinforcing effects of nicotine acting in worms were irrelevant to food. A development of CCP was also provoked by the repeated intermittent pairing of hexane with nicotine (Fig.4). CCP was successfully facilitated by the short period (1min) of multiple sessions of conditioning of US (nicotine) and CS (hexane) without *E. coli* (food) and following withdrawal, indicating CCP was specifically elicited by nicotine alone. It demonstrated the reinforcing effect of nicotine in *C. elegans*. Together with the result in Fig. 1d, in which CCP was not established when CS (hexane) was presented alone with the food, it demonstrated that nicotine is the primary reinforcer in the progress of CCP in *C. elegans*.

**Fig. 4.**
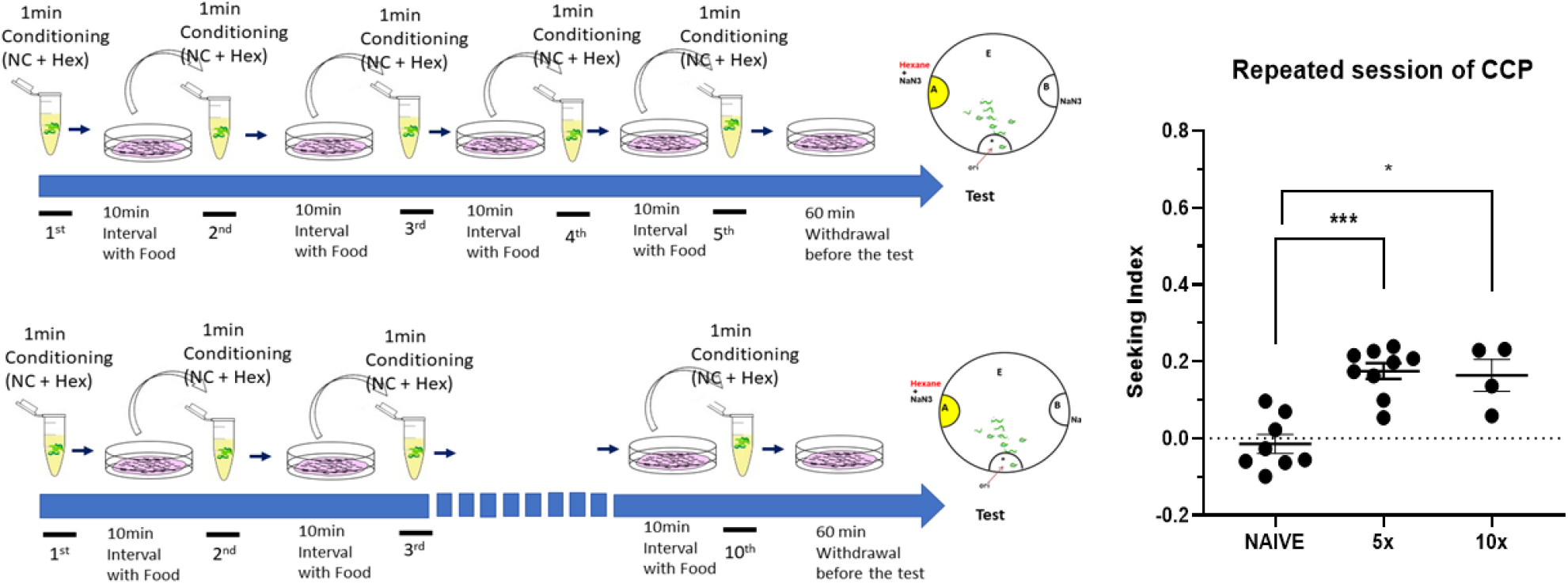
CCP is specifically elicited by nicotine. The short time of repeated conditioning (1min, without food during conditioning) and withdrawal elicits successful CCP. A conditioning session (nicotine and hexane) was 1minute and 10minutes of withdrawal was followed. After multiple session of conditioning, the last withdrawal session was consistent as 60minutes before conducting chemotaxis to CS. Each dot represents trial of population assay. *, P<0.05; ***, P<0.001 (Mann-Whitney test).

### CCP via nAChRs

The nAChRs function as pentameric ligand-gated ion channels. Conformational transitions after binding to nicotine, accompanied by various regulatory mechanisms enable nAChRs to respond dynamically to genetic and environmental factors. Elucidating subunits that specifically play a role in preference and seeking behavior elicited by nicotine will provide insight in the conservative role of nAChRs in mammals. It has been reported that two nAChR subunits, ACR-15 and ACR-16, are required for nicotine-withdrawal induced stimulation of locomotion ^23^. Nicotine also elicits associative learning with the rewarding effects in worm and ACR-5 and ACR-15 have been reported involved in this ^31^. We tested nAChR subunit KO mutant animals in CCP assay. The 29 nAChR homologs are reported in *C. elegans* genome whereas 17 in mammals ^23^,^43^. These nAChRs classified into five groups, which are ACR-16 group, UNC-29 group, UNC-38 group, ACR-8 group, and DEG-3 group ^43^. We screened 12 nAChR mutants by CCP assay, focusing on ACR-16 group, which closely resembles the mammalian α7-nAChR subunit, a predominant subtype in the brain ^44,45^. Here, we represent consistent results with previous findings and also newly identified additional nAChR subunits associated with nicotine-induced motivated behaviors (Fig. 5). In a single session of chronic CCP analysis, we identified delayed development of CCP in KO mutants of *acr-5*, and impaired in *acr-15, acr-16*, which is compatible with the previous reports in nicotine dependent-locomotion of worms (Fig. 5). Furthermore, we also identified the impaired development of CCP in KO mutants of *acr-9, acr-11, acr-21* (Fig. 5). The expression enrichment profile, provided by a single-cell gene expression profile of every neuron type in the *C. elegans* (CeNGEN) ^46^, shows that *acr-9* is expressed in AVA, a crucial interneuron validated for the development of nicotine-dependent locomotion, in which *acr-15* and *acr-16* are expressed^23^. Recently, the AVA interneurons have been shown to participate in the integration of sensory-motor input and decision making ^47^. Interestingly, *acr-21*, the nAChR α9 (CHRNA9) orthologue, is enriched in the RMG ^46^, the gap junctional hub interneurons that electrically connect to many sensory, motor, and interneurons and is known to modulate pheromone attraction and social behavior ^48^. RMG neurons form a close connection with AVA and ADA neurons, and *acr-11*, which we newly identified to play a role in nicotine CCP, is reported to be enriched in ADA. We also identified the *unc-63* and *unc-38* mutants were defective in the development of CCP. This result is consistent with previous investigation in nicotine dependent stimulation of locomotion, however, a further comprehensive analysis will be required as both mutant animals, *unc-63* and *unc-38*, are not severely uncoordinated as described ^23^,^49^,^50^. Nonetheless, our results demonstrate that the nicotine-elicited conditioned cue preference is mediated by nAChRs.

**Fig. 5.**
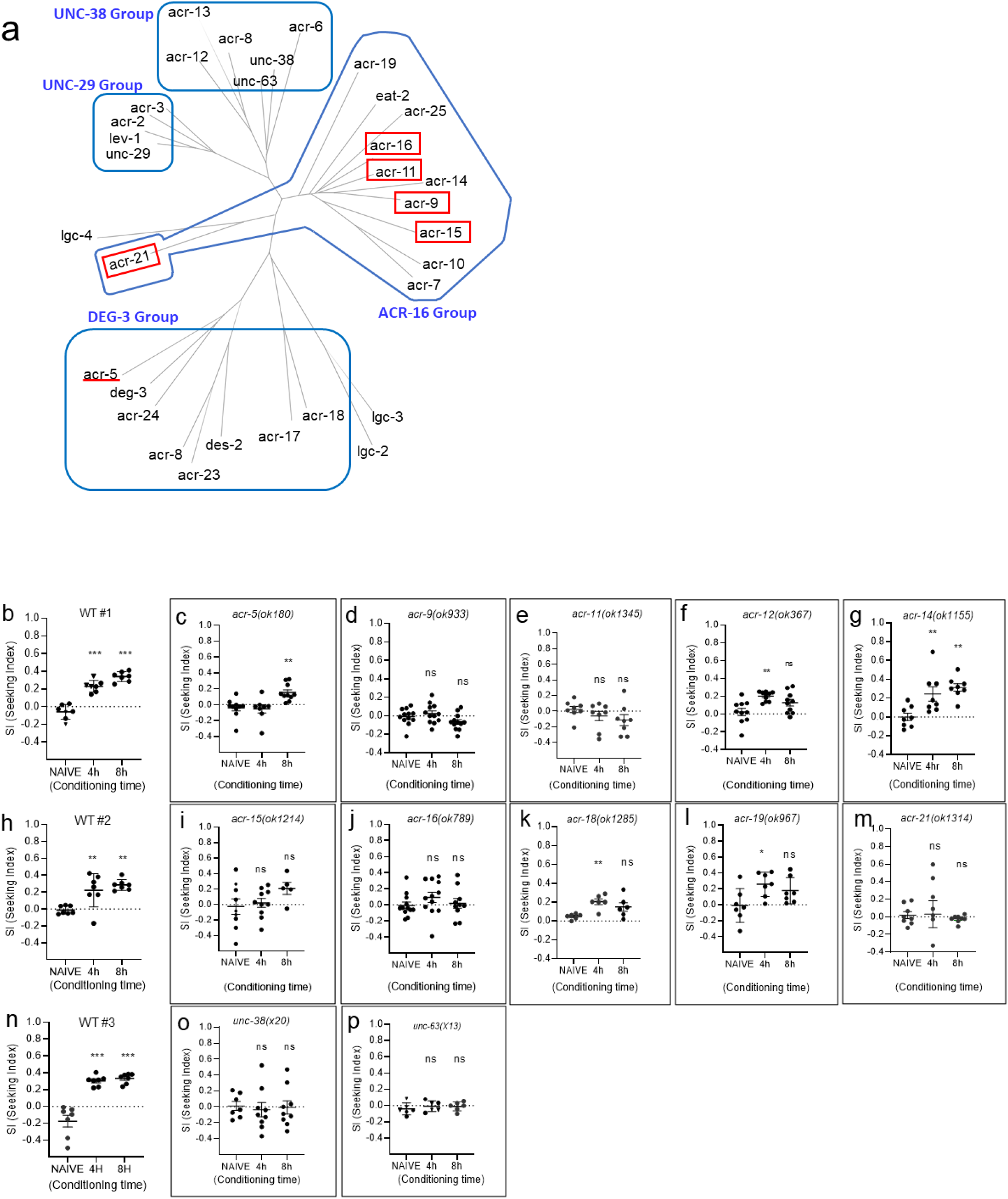
Identification of nAChRs relevant to CCP progression. (a) Phylogenetic analysis showing the nAChR receptor family of *C. elegans*. Using protein sequence homology, nAChR subunits were classified. (b) WT#1; one-way ANOVA with a post-hoc Dunnett’s test. F(2, 18)=62.13, P<0.001 (c) *acr-5 (ok180), (*P=0.002, One-way ANOVA, F(2, 23)=8.396, ** represents P<0.01 from post hoc multiple comparison test; Dunnett’s). ** represents P<0.01 from post hoc multiple comparison test; Dunnett’s. (d) *acr-9 (ok933)*, (<u>not significant</u>, One-way ANOVA, F(2, 33)=2.755, ns from post hoc multiple comparison test; Dunnett’s). (e) *acr-11(ok1345)*, (<u>not significant</u>, One-way ANOVA, F(2, 21)=1.423, ns from post hoc multiple comparison test; Dunnett’s). (f) *acr-12(ok367)*, (p=0.003, One-way ANOVA, F(2, 27)=7.029, ** represents P<0.01 from post hoc multiple comparison test; Dunnett’s). (g) *acr-14(ok1155)*, (p=0.001, One-way ANOVA, F(2, 21)=9.189, ** represents P<0.01 from post hoc multiple comparison test; Dunnett’s). (h) WT #2, (p<0.001, One-way ANOVA, F(2, 18)=11.17, ** represents P<0.01 from post hoc multiple comparison test; Dunnett’s). (i) *acr-15 (ok1214), (*<u>not significant</u>, One-way ANOVA, F(2, 21)=1.735, ns from post hoc multiple comparison test; Dunnett’s). (j) *acr-16 (ok789)*, (<u>not significant</u>, One-way ANOVA, F(2, 31)=0.9130, ns from post hoc multiple comparison test; Dunnett’s). (k) *acr-18(ok1285), (*P=0.01, One-way ANOVA, F(2, 15)=6.177, ** represents P<0.01 from post hoc multiple comparison test; Dunnett’s). (l) *acr-19(ok967)*, (P=0.03, One-way ANOVA, F(2, 18)=4.080, * represents P<0.05 from post hoc multiple comparison test; Dunnett’s). (m) *<u>acr-21(ok1314</u>)*, (<u>not significant</u>, One-way ANOVA, F(2, 21)=0.09735, ns from post hoc multiple comparison test; Dunnett’s). (n) WT#3, (p<0.001, One-way ANOVA, F(2, 18)=40.66, *** represents P<0.001 from post hoc multiple comparison test; Dunnett’s). (o) *unc-38(x20)*, (<u>not significant</u>, One-way ANOVA, F(2, 22)=0.08521, ns from post hoc multiple comparison test; Dunnett’s). (p) *unc-63(x13)*, (<u>not significant</u>, One-way ANOVA, F(2, 15)=0.5035, ns from post hoc multiple comparison test; Dunnett’s).

### Orthogonal test for nicotine preference

Cross-species functional validation of GWAS candidates using *C. elegans* has been used successfully to demonstrate the functional relevance of candidates in substance dependent behaviors (^51^). We asked whether nicotine CCP in worms could be a viable and useful tool to accelerate the assessment of biologically significant pathways associated with nicotine dependence through rapid functional characterization of GWAS candidates. Nicotine has been reported to evoke a calcium response from worms to mammals ^23,52-54^. The nAChRs mediate the increased intracellular calcium via VGCC-dependent and VGCC-independent manners that contribute to neural plasticity. Functional nAChRs are homopentameric or heteropentameric channels composed of 5 subunits by a combination of the α(α2-α10) and β(β2-β4) subunits ^55-58^. The Genome-wide meta-analysis on nicotine dependence has reported the protective role of *CACNA2D3* in nicotine dependence for African Americans ^59^. The *CACNA2D3* is also reported in the association of success in abstaining from smoking ^60^. *CACNA2D3* is responsible for encoding the α2δ, auxiliary subunits of Voltage-Gated Calcium Channel (VGCC), which influences the biophysical properties of the calcium channels ^61^. The worm orthologue of *CACNA2D3* modulates voltage dependence, the activation kinetics, and the conductance of calcium current of VGCC like mammalian a2δ (^62^). Other members of the α2δ family, *CACNA2D2* is also associated with nicotine dependence, smoking initiation, and cigarettes consumption ^63^. The loss of function alleles of *unc-36, CACNA2D3* orthologue was tested in CCP for the functional validation in the development of nicotine preference. We have tested multiple mutant alleles of *unc-36*. The *unc-36 (e251) and unc-36 (ad698)* are both loss of function alleles by the introduction of the premature stop codon and showed delayed or impaired progress of nicotine-conditioned cue preference in a single session of chronic CCP unlike WT animals (Fig. 6a, 6b, and 6c). We also tested mutant animals of *tag-180, CACNA2D2* orthologue, which is closely related to *CACNA2D3* in the same family and shared the human smoking phenotypes. The *tag-180 (ok779)*, deletion mutant (KO), showed impaired development of nicotine preference (Fig. 6d). We also tested animals in repeated session of conditioning and intermittent withdrawals. The orthogonal test exhibited a reduced development of CCP in unc-*36 (e251) and tag-180(ok779)* (Fig. 6e). Taken together, our data demonstrate that α2δ subunit of VGCC is required for the nicotine preference contributing to the development of nicotine dependence.

**Fig. 6.**
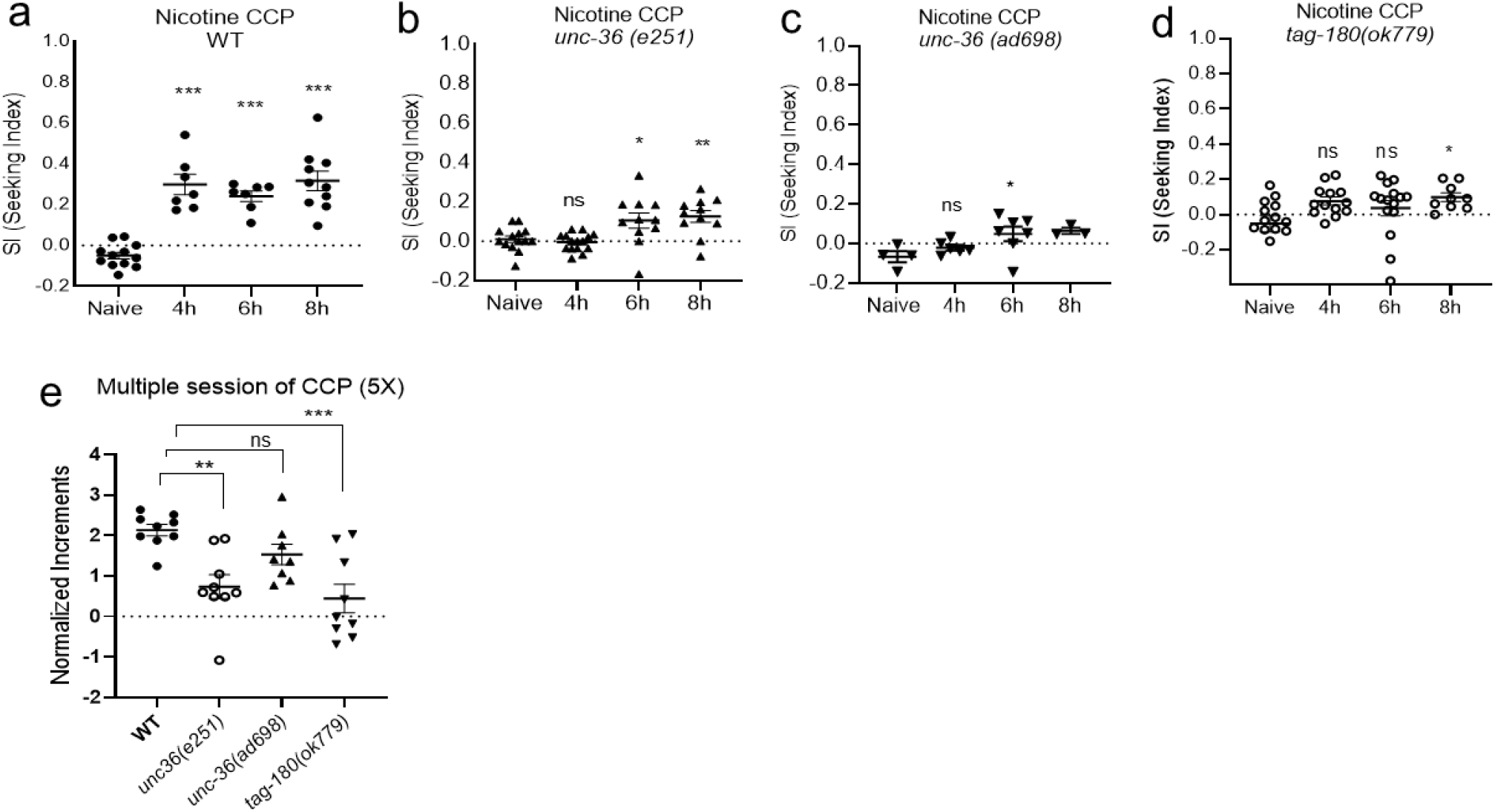
The orthogonal test evaluated nicotine preference of α2δ proteins (a) Wild-type CCP was conducted at each trial to evaluate the drug plate and the conditioning process (***, One-way ANOVA, F(3, 32)=27.17, *** represents P<0.001 from post hoc multiple comparison test; Dunnett’s). (b) *unc-36 (e251)*, orthologue of *CACNA2D3*, showed delayed and reduced development of CCP (***, One-way ANOVA, F(3, 47)=7.694, * represents p<0.05 and ** P<0.05 from post hoc multiple comparison test; Dunnett’s). (c) Impaired CCP was observed in *unc-36 (ad698)*, orthologue of *CACNA2D3*, (*, One-way ANOVA, F(3, 16)=3.475, * represents p<0.05 and ** P<0.05 from post hoc multiple comparison test; Dunnett’s). (d)Impaired CCP in *tag-180 (ok779)*, orthologue of *CACNA2D2*. (*, One-way ANOVA, F(3, 47)=2.885, * represents p<0.05 from post hoc multiple comparison test; Dunnett’s). (e) Orthogonal test in repeated CCP. Repeated training of conditioning and intermittent withdrawal further demonstrated reduced development of nicotine preference in the mutant animals of α2δ orthologues. (***, One-way ANOVA, F(3, 31)=8.112, ** represents p<0.01 and *** P<0.001 from post hoc multiple comparison test; Dunnett’s). Each dot represents a trial of population assay.

## Discussion

Human genetic association studies have been successful in revealing genetic variants associated with smoking-related phenotypes. When analyzing the NHGRI/EBI GWAS catalog (release:2021-01-14), it contained 1,504 SNPs associated with smoking/nicotine that reached genome-wide significance. Most of these variants (93%) are yet to be replicated by an independent study. Although GWAS studies associated with tobacco smoking have revealed numerous genetic factors, the estimated heritability has been limited to explaining underlying mechanisms. Thus, various attempts have been suggested to accelerate functional validation and comprehensive analysis ^64^. In a comparative proteomics study, 83% of the worm proteome exhibits homology with human genes and recent meta-analysis with orthology-prediction methods showed that approximately 52.6% of the human protein-coding genome has noticeable orthologues in worms, illustrating that the nematode provides a suitable model organism for functional validation of human genes. ^65,66^.

Cross-species functional validation has long been used in worms including SUD related phenotypes. For example, the introduction of a human TRPC (transient receptor potential canonical) channel can rescue the defective nicotine dependent simulated locomotion phenotype of worm TRPC channel KO strain ^23^. A mammalian transient receptor potential channel vanilloid (TRPV) can substitute worm orthologue and directs behavioral responses ^67^. The transgenic worms containing the human SLC18A2 gene provided model to investigate the brain dopamine and serotonin vesicular transport disease ^68^. Recently, interspecies chimerism with a mammalian gene in the worm platform identified orphan anti-opioid system ^69^. The transgenic worm to express the mammalian μ (mu) opioid receptor (MOR), which is not normally found in the worm genome, responds to opioids such as morphine and fentanyl. Successively, this transgenic worm contributed to finding the orphan GPCR of which mammalian orthologue shows functional conservation related to the anti-opioid pathway. Therefore, we exploited worms to define vulnerability phenotypes by proper modeling of behavioral phenotypes and to test the functional evaluation of human GWAS candidates associated with nicotine dependence and smoking. The CCP in worms is specifically induced by nicotine and mediated by dopamine. We identified the nicotine-elicited cue preference is mediated by nicotinic acetylcholine receptors in worms. Taken together, we demonstrated that worms exhibited the key features of nicotine-dependent behaviors in mammals.

The GWAS reveals numerous risk factors associated with diseases. Despite the successes of GWAS, most of the candidate genetic variants have not been independently validated or provided novel insight into novel treatments. Experimental approaches for functional validation will be required to determine whether candidate genes have an actual role in the disease. A previously identified GWAS variant of *CACNA2D3*, in which the SNP is in the intron region, was not prioritized for further validation, but it was reported that this variant was associated with reduced expression levels in three human brain tissues and was associated with nicotine dependence ^59^. We validated its function by testing the loss of function or KO strains of orthologue that allow for further pathway evaluation afterwards. A *CACNA2D3* encodes α2δ, auxiliary subunits of VGCC, that influences the biophysical properties of the calcium channels. VGCCs are pivotal in excitable cells with permeability to mainly calcium ions. Although it has been suggested that permeation of calcium ions into cells through VGCC will play a pivotal role in the induction of plasticity of nicotine through nAChRs ^52,70,71^, close interaction between nAChRs and VGCC for the subsequent event to mediate nicotine response is depending on the cell types, in which specific subtypes of nAChRs are expressed ^57,58^. Mostly, non-α7-nAChRs mainly interact with the VGCC to mediate the signaling caused by nicotine.

α2δ proteins are encoded by 4 genes (*CACNA2D1, CACNA2D2, CACNA2D3, CACNA2D4*) and expressed throughout the central nervous system to co-assemble with most of the α1 subunit forming functional calcium channel ^72^. α2δ proteins also interact with other proteins such as α-neurexins, LRP1 (low-density lipoprotein receptor-related protein 1), NMDA receptor (N-methyl-d-aspartate), and BK channels (large-conductance calcium-activated potassium channels) ^73-76^. The part of these might be related to recent implications of α2δ proteins in the progress of SUD. Like *CACNA2D2* and *CACNA2D3* have been reported as GWAS candidates associated with nicotine dependence ^59,63^, CACNA2D1 has been involved in the increased presynaptic NMDAR activity associated with hyperalgesia following chronic morphine ^77^. An aberrant interaction between thrombospondin (TSP) and CACNA2D1 has been proposed as a possible mechanism of synaptic remodeling in the hippocampus during chronic ethanol consumption ^78^. An interaction between α-neurexins and α2δ proteins is evolutionarily well conserved endorsed by an interaction between NRX-1 and UNC-36 in *C. eleagns* ^73^. The *C. elegans* genome includes 2 genes predicted to encode α2δ family proteins, *unc-36* and *tag-180*, predicted as *CACNA2D3/ CACNA2D1* like orthologue and *CACNA2D2* like orthologue, respectively ^66^. Like mammalian α2δ proteins, the function of UNC-36 in the modulation of the voltage dependence, the activation kinetics, and the conductance of calcium currents was electrophysiologically validated in the neuromuscular junction, whereas TAG-180 has no effects ^62^. UNC-36 has been also demonstrated as a regulator of synaptogenesis together with UNC-2, Ca_v_2-like α1 subunit of VGCC, in the neuromuscular junction ^79^. Interestingly, the *tag-180* has not shown a functional association related to calcium channel activity so far. However, it is of interest that the behavioral phenotype of *tag-180* in nicotine motivated behavior has been defined, here. Perhaps it reflects the non-canonical interactions and role of α2δ proteins, such as the accumulation of CACNA2D2 in lipid rafts independently from the interaction with calcium channels ^80^.

Here, we have established a novel CPP paradigm assay for nicotine seeking in worms which could accelerate the functional validation of genes associated with the progress of nicotine dependence. We determined the nicotine seeking by CCP and the functional validation of the orthogonal test showed orthologues of *CACNA2D2, CACNA2D3* have a role in nicotine-motivated behavior in *C. elegans*. Thus, follow-up studies of the α2δ protein should be performed to investigate a comprehensive functional characterization of the mechanisms of nicotine seeking and taking. We are pursuing the identification of nAChR subunits that specifically act on nicotine seeking and defining a subset of neurons in which this subunit acts.

## Materials and Methods

All strains were cultivated on nematode growth media (NGM) plates with *Escherichia coli* strain OP50 at 20°C as described ^81^ and the hermaphrodite worm was used for behavioral analysis. The Bristol N2 strain was used as wild-type (WT) animals. The strains below were obtained from Caenorhabditis Genetics Center (CGC, Minneapolis, MN, USA), which is supported by the National Institutes of Health Office of Research Infrastructure Programs (P40 OD010440). The following mutant alleles were used in the study: *cat-2 (e1112), acr-5(ok180), acr-9(ok933), acr-11(ok1345), acr-12(ok367), acr-14(ok1155), acr-15(ok1214), acr-16(ok789), acr-18(ok1285), acr-19(ok967), acr-21(ok1314), unc-38 (x20), and unc-63(x13), unc-36(e251), unc-36(ad698), tag-180(ok779)*. The strain PY7502, oyIs85[*ceh-36p*::TU#813 + *ceh-36*::TU#814 + *srtx-1p*::GFP + *unc122p*::DsRed], was used for AWC ablated animals. PY7502 was generated via expression of recCaspases (split caspases) ^41^ under *ceh-36* promoter ^82^.

### Behavioral Assay

#### Nicotine Conditioning

The nicotine plates were prepared freshly in 60 mm plate. When NGM is cooled to 55°C after sterilization, nicotine was added up to the designated concentration (1.5 μM). Concentrated OP50 was seeded on the nicotine plates and then one day later nicotine plates were used for conditioning. OP50-seeded nicotine plates were stored at 4°C and consumed within a week for the conditioning.

The Synchronized eggs were collected for 3 hours, and then were harvested with S basal-buffer [100 mM sodium chloride, 50 mM potassium phosphate (pH 6.0)] for the conditioning when they were reached to Day1 young adult stage (16-24 hours later after mid-L4 stage). To introduce hexane as a Conditioned Stimulus (CS) to the nicotine conditioning plate, 80 µl of agar lump (2% BBL agar) on the lid (60 mm plate) was freshly prepared before the conditioning. S basal-buffer harvested animals were placed in the middle of the conditioning plate (1.5 μM nicotine) and then covered with a lid with an agar lump which 3µl of hexane was added. Since a CS was a volatile odor, the plates were sealed with parafilm during conditioning. After 4, 6, or 8 hours of conditioning (for 8hrs, 3 µl of hexane was refilled to agar lump after 4hrs for another 4hr), worms were washed with S basal -buffer 3 times then transferred to OP50 seeded NGM without nicotine and hexane for the withdrawal session. 1 hour later, a chemotaxis assay was conducted. The withdrawal procedure (here, 1 hour) was followed after all the sessions, including [CS only] and [US only] which validated CCP, prior to performing chemotaxis to CS.

For the repeated sessions of conditioning, 1 minute of conditioning was conducted in 1ml of the S basal -buffer containing 1.5 μM nicotine and 2 μl of hexane with gentle rotating. After washing with S basal 3 times, conditioned worms were placed on OP50-seeded NGM for 10 minutes of withdrawal session. The above was performed repeatedly. The final withdrawal session prior to performing chemotaxis to CS was 1 hour like other CCP.

#### Chemotaxis to CS

A chemotaxis assay was performed as described previously ^36,39,83^. Briefly, 10 ml chemotaxis media [1.6% BBL-agar, 5mM potassium phosphate; pH 6.0, 1mM CaCl_2_, 1mM MgSO_4_] were prepared on the 100 mm petri dish. 1 µl of 100mM NaN_3_ was added to the point marked in the section of A and B (Fig. 1b). 1 µl of CS (undiluted hexane) was added on top of the NaN_3_ in the section of A. Immediately after the CS was absorbed into the 100 mm chemotaxis plate, about 100 washed animals were placed in the area marked using glassware micropipette. 40 minutes later with parafilm sealing, the number of accumulated animals in each section marked (Fig. 1b) was counted to calculate the Seeking index. The index was calculated by [(number of animals in A - number of animals in B)/Total number of animals [Seeking index SI= (A-B)/Total(A+B+E)]. Total 100-150 animals were tested in each trial to get the index.

In the case of an uncoordinated strains [*unc-38 (x20), and unc-63(x13), unc-36(e251), unc-36(ad698)*], their CCP was confirmed again by creating an environment that could be reached to the CS (same concentration given) by moving a short distance. A square 100 mm chemotaxis plate with a grid engraved on it was prepared using the same amount of chemotaxis media. And then chemotaxis was performed in a space where animals showing uncoordinated movement using only 60 mm in the center could arrive at their destination in time. At these trials, the WT control were also performed under the same conditions.

### Statistical Analysis

WT control groups were always tested together at each trial to evaluate the drug plate and the conditioning process. Each dot in the graph represents the population assay in which about 100-150 animals were tested. The mean and standard error of the mean (SEM) were determined for all experimental parameters. The data were analyzed employing the Mann-Whitney or Dunnett’s tests using GraphPad Prism software (version 8.0.1). Data points with p-values below 0.05 (P < 0.05) were considered to be significant.

### Sequence alignment

protein sequences were analyzed by database similarity search (^84^) and the multiple protein sequences were simultaneously aligned using the COBALT, a constraint based alignment tool (^85^). The phylogenetic tree was constructed by COBALT using minimum evolution method. The sequences used for the phylogenetic tree analysis: P48182.1, Q93149.1, P54246.5, NP_491354.2, NP_495647.1, NP_001361818.1, NP_510285.2, NP_508692.3, NP_491906.1, AAG35183.1, NP_495716.1, NP_505206.2, NP_505207.1, NP_001023961.1, NP_506868.2, NP_001129756.1, NP_001367183.1, NP_001355515.1, NP_504024.2, NP_001379138.1, NP_001380111.1, NP_496959.1, NP_001255705.1, NP_509932.2, NP_492399.1, NP_491472.1, NP_491533.2, G5ECT0.1, NP_001255865.1, Q19351.5, NP_509556.4, NP_001023570.2

## Acknowledgments

This work was supported by College of Medicine, University of Tennessee Health Science Center (UTHSC) and a pilot project in support of P50DA037844, University of California San Diego. We thank the *C. elegans* Genetics Center (CGC) for providing strains, which is funded by NIH Office of Research Infrastructure Programs (P40 OD010440). We also thank the CeNGEN(a single-cell gene expression profile of every neuron type in the *C. elegans*: Funded by NIH-NINDS, grant R01NS100547) for providing gene expression profile. Authors declare no conflict of interests.

## References

1 WorldHealthOrganization. WHO Report on the Global Tobacco Epidemic. 2017external icon, Geneva (2017).

2 Koopmans, J. R., Slutske, W. S., Heath, A. C., Neale, M. C. & Boomsma, D. I. The genetics of smoking initiation and quantity smoked in Dutch adolescent and young adult twins. Behav Genet 29, 383–393, doi:10.1023/a:1021618719735 (1999).

3 Stallings, M. C., Hewitt, J. K., Beresford, T., Heath, A. C. & Eaves, L. J. A twin study of drinking and smoking onset and latencies from first use to regular use. Behav Genet 29, 409–421, doi:10.1023/a:1021622820644 (1999).

4 Heath, A. C., Kirk, K. M., Meyer, J. M. & Martin, N. G. Genetic and social determinants of initiation and age at onset of smoking in Australian twins. Behav Genet 29, 395–407, doi:10.1023/a:1021670703806 (1999).

5 Vink, J. M., Willemsen, G. & Boomsma, D. I. Heritability of smoking initiation and nicotine dependence. Behav Genet 35, 397–406, doi:10.1007/s10519-004-1327-8 (2005).

6 Hall, W. D., Gartner, C. E. & Carter, A. The genetics of nicotine addiction liability: ethical and social policy implications. Addiction 103, 350–359, doi:10.1111/j.1360-0443.2007.02070.x (2008).

7 Hancock, D. B. et al. Genome-wide meta-analysis reveals common splice site acceptor variant in CHRNA4 associated with nicotine dependence. Transl Psychiatry 5, e651, doi:10.1038/tp.2015.149 (2015).

8 Gorwood, P., Le Strat, Y. & Ramoz, N. Genetics of addictive behavior: the example of nicotine dependence. Dialogues Clin Neurosci 19, 237–245 (2017).

9 Gelernter, J. et al. Genome-wide association study of nicotine dependence in American populations: identification of novel risk loci in both African-Americans and European-Americans. Biol Psychiatry 77, 493–503, doi:10.1016/j.biopsych.2014.08.025 (2015).

10 Gelernter, J. et al. Genome-wide association study of opioid dependence: multiple associations mapped to calcium and potassium pathways. Biol Psychiatry 76, 66–74, doi:10.1016/j.biopsych.2013.08.034 (2014).

11 Gelernter, J. et al. Genome-wide association study of cocaine dependence and related traits: FAM53B identified as a risk gene. Mol Psychiatry 19, 717–723, doi:10.1038/mp.2013.99 (2014).

12 Gelernter, J. et al. Genome-wide association study of alcohol dependence:significant findings in African- and European-Americans including novel risk loci. Mol Psychiatry 19, 41–49, doi:10.1038/mp.2013.145 (2014).

13 Klein, R. J., Xu, X., Mukherjee, S., Willis, J. & Hayes, J. Successes of genome-wide association studies. Cell 142, 350-351; author reply 353-355, doi:10.1016/j.cell.2010.07.026 (2010).

14 Hirschhorn, J. N. Genomewide association studies--illuminating biologic pathways. N Engl J Med 360, 1699–1701, doi:10.1056/NEJMp0808934 (2009).

15 White, J. G., Southgate, E., Thomson, J. N. & Brenner, S. The structure of the nervous system of the nematode Caenorhabditis elegans. Philosophical transactions of the Royal Society of London. Series B, Biological sciences 314, 1–340, doi:10.1098/rstb.1986.0056 (1986).

16 Varshney, L. R., Chen, B. L., Paniagua, E., Hall, D. H. & Chklovskii, D. B. Structural properties of the Caenorhabditis elegans neuronal network. PLoS computational biology 7, doi:10.1371/journal.pcbi.1001066 (2011).

17 Jarrell, T. A. et al. The connectome of a decision-making neural network. Science (New York, N.Y.) 337, 437–444, doi:10.1126/science.1221762 (2012).

18 Jee, C. et al. SEB-3, a CRF receptor-like GPCR, regulates locomotor activity states, stress responses and ethanol tolerance in Caenorhabditis elegans. Genes, brain, and behavior 12, 250–262, doi:10.1111/j.1601-183X.2012.00829.x (2013).

19 Bierut, L. J. Genetic vulnerability and susceptibility to substance dependence. Neuron 69, 618–627, doi:10.1016/j.neuron.2011.02.015 (2011).

20 Davies, A. G. et al. A central role of the BK potassium channel in behavioral responses to ethanol in C. elegans. Cell 115, 655–666, doi:10.1016/S0092-8674(03)00979-6 (2003).

21 Davies, A. G., Bettinger, J. C., Thiele, T. R., Judy, M. E. & McIntire, S. L. Natural variation in the npr-1 gene modifies ethanol responses of wild strains of C. elegans. Neuron 42, 731–743, doi:10.1016/j.neuron.2004.05.004 (2004).

22 Lee, J., Jee, C. & McIntire, S. L. Ethanol preference in C. elegans. Genes, brain, and behavior 8, 578–585, doi:10.1111/j.1601-183X.2009.00513.x (2009).

23 Feng, Z. et al. A C. elegans model of nicotine-dependent behavior: regulation by TRP-family channels. Cell 127, 621–633, doi:10.1016/j.cell.2006.09.035 (2006).

24 Ward, A., Walker, V. J., Feng, Z. & Xu, X. Z. Cocaine modulates locomotion behavior in C. elegans. PloS one 4, doi:10.1371/journal.pone.0005946 (2009).

25 Carvelli, L., Matthies, D. S. & Galli, A. Molecular mechanisms of amphetamine actions in Caenorhabditis elegans. Molecular pharmacology 78, 151–156, doi:10.1124/mol.109.062703 (2010).

26 Waggoner, L. E. et al. Long-term nicotine adaptation in Caenorhabditis elegans involves PKC-dependent changes in nicotinic receptor abundance. The Journal of neuroscience : the official journal of the Society for Neuroscience 20, 8802–8811 (2000).

27 Rauthan, M. et al. MicroRNA Regulation of nAChR Expression and Nicotine-Dependent Behavior in C. elegans. Cell Reports 21, 1434–1441, doi:10.1016/j.celrep.2017.10.043 (2017).

28 Fudala, P. J., Teoh, K. W. & Iwamoto, E. T. Pharmacologic characterization of nicotine-induced conditioned place preference. Pharmacol Biochem Behav 22, 237–241, doi:10.1016/0091-3057(85)90384-3 (1985).

29 Spyraki, C., Fibiger, H. C. & Phillips, A. G. Dopaminergic substrates of amphetamine-induced place preference conditioning. Brain Res 253, 185–193, doi:10.1016/0006-8993(82)90685-0 (1982).

30 Kruszewska, A., Romandini, S. & Samanin, R. Different effects of zimelidine on the reinforcing properties of d-amphetamine and morphine on conditioned place preference in rats. Eur J Pharmacol 125, 283–286, doi:10.1016/0014-2999(86)90038-5 (1986).

31 Sellings, L. et al. Nicotine-motivated behavior in Caenorhabditis elegans requires the nicotinic acetylcholine receptor subunits acr-5 and acr-15. The European journal of neuroscience 37, 743–756, doi:10.1111/ejn.12099 (2013).

32 Di Chiara, G. Role of dopamine in the behavioural actions of nicotine related to addiction. Eur J Pharmacol 393, 295–314, doi:10.1016/s0014-2999(00)00122-9 (2000).

33 Ikemoto, S., Qin, M. & Liu, Z. H. Primary reinforcing effects of nicotine are triggered from multiple regions both inside and outside the ventral tegmental area. J Neurosci 26, 723–730, doi:10.1523/jneurosci.4542-05.2006 (2006).

34 Wes, P. D. & Bargmann, C. I. C. elegans odour discrimination requires asymmetric diversity in olfactory neurons. Nature 410, 698–701, doi:10.1038/35070581 (2001).

35 Hukema, R. K., Rademakers, S. & Jansen, G. Gustatory plasticity in C. elegans involves integration of negative cues and NaCl taste mediated by serotonin, dopamine, and glutamate. Learning & memory (Cold Spring Harbor, N.Y.) 15, 829–836, doi:10.1101/lm.994408 (2008).

36 Bargmann, C. I., Hartwieg, E. & Horvitz, H. R. Odorant-selective genes and neurons mediate olfaction in C. elegans. Cell 74, 515–527, doi:10.1016/0092-8674(93)80053-h (1993).

37 Hilliard, M. A., Bergamasco, C., Arbucci, S., Plasterk, R. H. & Bazzicalupo, P. Worms taste bitter: ASH neurons, QUI-1, GPA-3 and ODR-3 mediate quinine avoidance in Caenorhabditis elegans. The EMBO journal 23, 1101–1111, doi:10.1038/sj.emboj.7600107 (2004).

38 Sengupta, P., Colbert, H. A. & Bargmann, C. I. The C. elegans gene odr-7 encodes an olfactory-specific member of the nuclear receptor superfamily. Cell 79, 971–980, doi:10.1016/0092-8674(94)90028-0 (1994).

39 Bargmann, C. I. & Horvitz, H. R. Chemosensory neurons with overlapping functions direct chemotaxis to multiple chemicals in C. elegans. Neuron 7, 729–742, doi:10.1016/0896-6273(91)90276-6 (1991).

40 Larsch, J. et al. A Circuit for Gradient Climbing in C. elegans Chemotaxis. Cell reports 12, 1748–1760, doi:10.1016/j.celrep.2015.08.032 (2015).

41 Chelur, D. S. & Chalfie, M. Targeted cell killing by reconstituted caspases. Proceedings of the National Academy of Sciences of the United States of America 104, 2283–2288, doi:10.1073/pnas.0610877104 (2007).

42 Beverly, M., Anbil, S. & Sengupta, P. Degeneracy and neuromodulation among thermosensory neurons contribute to robust thermosensory behaviors in Caenorhabditis elegans. The Journal of neuroscience : the official journal of the Society for Neuroscience 31, 11718–11727, doi:10.1523/JNEUROSCI.1098-11.2011 (2011).

43 Polli, J. R. et al. Drug-dependent behaviors and nicotinic acetylcholine receptor expressions in Caenorhabditis elegans following chronic nicotine exposure. NeuroToxicology 47, 27–36, doi:10.1016/j.neuro.2014.12.005 (2015).

44 Brejc, K. et al. Crystal structure of an ACh-binding protein reveals the ligand-binding domain of nicotinic receptors. Nature 411, 269–276, doi:10.1038/35077011 (2001).

45 Sharma, G. & Vijayaraghavan, S. Nicotinic receptors containing the alpha7 subunit: a model for rational drug design. Curr Med Chem 15, 2921–2932, doi:10.2174/092986708786848703 (2008).

46 Hammarlund, M., Hobert, O., Miller, D. M., 3rd & Sestan, N. The CeNGEN Project: The Complete Gene Expression Map of an Entire Nervous System. Neuron 99, 430–433, doi:10.1016/j.neuron.2018.07.042 (2018).

47 Liu, P., Chen, B. & Wang, Z. W. GABAergic motor neurons bias locomotor decision-making in C. elegans. Nat Commun 11, 5076, doi:10.1038/s41467-020-18893-9 (2020).

48 Macosko, E. Z. et al. A hub-and-spoke circuit drives pheromone attraction and social behaviour in C. elegans. Nature 458, 1171–1175, doi:10.1038/nature07886 (2009).

49 Fleming, J. T. et al. Caenorhabditis elegans levamisole resistance genes lev-1, unc-29, and unc-38 encode functional nicotinic acetylcholine receptor subunits. J Neurosci 17, 5843–5857, doi:10.1523/jneurosci.17-15-05843.1997 (1997).

50 Culetto, E. et al. The Caenorhabditis elegans unc-63 gene encodes a levamisole-sensitive nicotinic acetylcholine receptor alpha subunit. J Biol Chem 279, 42476–42483, doi:10.1074/jbc.M404370200 (2004).

51 Adkins, A. E. et al. Genomewide Association Study of Alcohol Dependence Identifies Risk Loci Altering Ethanol-Response Behaviors in Model Organisms. Alcohol Clin Exp Res 41, 911–928, doi:10.1111/acer.13362 (2017).

52 Campusano, J. M., Su, H., Jiang, S. A., Sicaeros, B. & O’Dowd, D. K. nAChR-mediated calcium responses and plasticity in Drosophila Kenyon cells. Dev Neurobiol 67, 1520–1532, doi:10.1002/dneu.20527 (2007).

53 Vijayaraghavan, S., Pugh, P. C., Zhang, Z. W., Rathouz, M. M. & Berg, D. K. Nicotinic receptors that bind alpha-bungarotoxin on neurons raise intracellular free Ca2+. Neuron 8, 353–362, doi:10.1016/0896-6273(92)90301-s (1992).

54 Sharma, G., Grybko, M. & Vijayaraghavan, S. Action potential-independent and nicotinic receptor-mediated concerted release of multiple quanta at hippocampal CA3-mossy fiber synapses. J Neurosci 28, 2563–2575, doi:10.1523/jneurosci.5407-07.2008 (2008).

55 Albuquerque, E. X. et al. Properties of neuronal nicotinic acetylcholine receptors: pharmacological characterization and modulation of synaptic function. J Pharmacol Exp Ther 280, 1117–1136 (1997).

56 Changeux, J. P. Nicotine addiction and nicotinic receptors: lessons from genetically modified mice. Nat Rev Neurosci 11, 389–401, doi:10.1038/nrn2849 (2010).

57 Dajas-Bailador, F. A., Mogg, A. J. & Wonnacott, S. Intracellular Ca2+ signals evoked by stimulation of nicotinic acetylcholine receptors in SH-SY5Y cells: contribution of voltage-operated Ca2+ channels and Ca2+ stores. J Neurochem 81, 606–614, doi:10.1046/j.1471-4159.2002.00846.x (2002).

58 Dajas-Bailador, F. & Wonnacott, S. Nicotinic acetylcholine receptors and the regulation of neuronal signalling. Trends Pharmacol Sci 25, 317–324, doi:10.1016/j.tips.2004.04.006 (2004).

59 Yin, X. et al. Genome-wide meta-analysis identifies a novel susceptibility signal at CACNA2D3 for nicotine dependence. Am J Med Genet B Neuropsychiatr Genet 174, 557–567, doi:10.1002/ajmg.b.32540 (2017).

60 Uhl, G. R. et al. Molecular genetics of successful smoking cessation: convergent genome-wide association study results. Arch Gen Psychiatry 65, 683–693, doi:10.1001/archpsyc.65.6.683 (2008).

61 Davies, A. et al. Functional biology of the alpha(2)delta subunits of voltage-gated calcium channels. Trends Pharmacol Sci 28, 220–228, doi:10.1016/j.tips.2007.03.005 (2007).

62 Lainé, V., Frøkjær-Jensen, C., Couchoux, H. & Jospin, M. The alpha1 subunit EGL-19, the alpha2/delta subunit UNC-36, and the beta subunit CCB-1 underlie voltage-dependent calcium currents in Caenorhabditis elegans striated muscle. J Biol Chem 286, 36180–36187, doi:10.1074/jbc.M111.256149 (2011).

63 Liu, M. et al. Association studies of up to 1.2 million individuals yield new insights into the genetic etiology of tobacco and alcohol use. Nat Genet 51, 237–244, doi:10.1038/s41588-018-0307-5 (2019).

64 Xu, Y. et al. Prediction of Smoking Behavior From Single Nucleotide Polymorphisms With Machine Learning Approaches. Front Psychiatry 11, 416, doi:10.3389/fpsyt.2020.00416 (2020).

65 Lai, C. H., Chou, C. Y., Ch’ang, L. Y., Liu, C. S. & Lin, W. Identification of novel human genes evolutionarily conserved in Caenorhabditis elegans by comparative proteomics. Genome Res 10, 703–713, doi:10.1101/gr.10.5.703 (2000).

66 Kim, W., Underwood, R. S., Greenwald, I. & Shaye, D. D. OrthoList 2: A New Comparative Genomic Analysis of Human and Caenorhabditis elegans Genes. Genetics 210, 445–461, doi:10.1534/genetics.118.301307 (2018).

67 Liedtke, W., Tobin, D. M., Bargmann, C. I. & Friedman, J. M. Mammalian TRPV4 (VR-OAC) directs behavioral responses to osmotic and mechanical stimuli in Caenorhabditis elegans. Proc Natl Acad Sci U S A 100 Suppl 2, 14531–14536, doi:10.1073/pnas.2235619100 (2003).

68 Young, A. T. et al. Modelling brain dopamine-serotonin vesicular transport disease in Caenorhabditis elegans. Dis Model Mech 11, doi:10.1242/dmm.035709 (2018).

69 Wang, D. et al. Genetic behavioral screen identifies an orphan anti-opioid system. Science 365, 1267–1273, doi:10.1126/science.aau2078 (2019).

70 Katsura, M. et al. Up-regulation of L-type voltage-dependent calcium channels after long term exposure to nicotine in cerebral cortical neurons. J Biol Chem 277, 7979–7988, doi:10.1074/jbc.M109466200 (2002).

71 Michalak, A. & Biala, G. Calcium homeostasis and protein kinase/phosphatase balance participate in nicotine-induced memory improvement in passive avoidance task in mice. Behav Brain Res 317, 27–36, doi:10.1016/j.bbr.2016.09.023 (2017).

72 Risher, W. C. & Eroglu, C. Emerging roles for α2δ subunits in calcium channel function and synaptic connectivity. Curr Opin Neurobiol 63, 162–169, doi:10.1016/j.conb.2020.04.007 (2020).

73 Tong, X. J. et al. Retrograde Synaptic Inhibition Is Mediated by α-Neurexin Binding to the α2δ Subunits of N-Type Calcium Channels. Neuron 95, 326-340.e325, doi:10.1016/j.neuron.2017.06.018 (2017).

74 Chen, J. et al. The α2δ-1-NMDA Receptor Complex Is Critically Involved in Neuropathic Pain Development and Gabapentin Therapeutic Actions. Cell Rep 22, 2307–2321, doi:10.1016/j.celrep.2018.02.021 (2018).

75 Zhang, F. X., Gadotti, V. M., Souza, I. A., Chen, L. & Zamponi, G. W. BK Potassium Channels Suppress Cavα2δ Subunit Function to Reduce Inflammatory and Neuropathic Pain. Cell Rep 22, 1956–1964, doi:10.1016/j.celrep.2018.01.073 (2018).

76 Kadurin, I., Rothwell, S. W., Lana, B., Nieto-Rostro, M. & Dolphin, A. C. LRP1 influences trafficking of N-type calcium channels via interaction with the auxiliary α(2)δ-1 subunit. Scientific reports 7, 43802–43802, doi:10.1038/srep43802 (2017).

77 Deng, M., Chen, S. R., Chen, H. & Pan, H. L. α2δ-1-Bound N-Methyl-D-aspartate Receptors Mediate Morphine-induced Hyperalgesia and Analgesic Tolerance by Potentiating Glutamatergic Input in Rodents. Anesthesiology 130, 804–819, doi:10.1097/aln.0000000000002648 (2019).

78 Risher, M. L. et al. Adolescent Intermittent Alcohol Exposure: Dysregulation of Thrombospondins and Synapse Formation are Associated with Decreased Neuronal Density in the Adult Hippocampus. Alcohol Clin Exp Res 39, 2403–2413, doi:10.1111/acer.12913 (2015).

79 Caylor, R. C., Jin, Y. & Ackley, B. D. The Caenorhabditis elegans voltage-gated calcium channel subunits UNC-2 and UNC-36 and the calcium-dependent kinase UNC-43/CaMKII regulate neuromuscular junction morphology. Neural Dev 8, 10, doi:10.1186/1749-8104-8-10 (2013).

80 Davies, A. et al. The calcium channel alpha2delta-2 subunit partitions with CaV2.1 into lipid rafts in cerebellum: implications for localization and function. J Neurosci 26, 8748–8757, doi:10.1523/jneurosci.2764-06.2006 (2006).

81 Brenner, S. The genetics of Caenorhabditis elegans. Genetics 77, 71–94 (1974).

82 Kim, K., Kim, R. & Sengupta, P. The HMX/NKX homeodomain protein MLS-2 specifies the identity of the AWC sensory neuron type via regulation of the ceh-36 Otx gene in C. elegans. Development (Cambridge, England) 137, 963–974, doi:10.1242/dev.044719 (2010).

83 Colbert, H. A. & Bargmann, C. I. Odorant-specific adaptation pathways generate olfactory plasticity in C. elegans. Neuron 14, 803–812, doi:10.1016/0896-6273(95)90224-4 (1995).

84 Gish, W. & States, D. J. Identification of protein coding regions by database similarity search. Nat Genet 3, 266–272, doi:10.1038/ng0393-266 (1993).

85 Papadopoulos, J. S. & Agarwala, R. COBALT: constraint-based alignment tool for multiple protein sequences. Bioinformatics 23, 1073–1079, doi:10.1093/bioinformatics/btm076 (2007).

